# Both Determinants of Allosteric and Active Sites Responsible for Catalytic Activity of Delta 12 Fatty Acid Desaturase by Domain Swapping

**DOI:** 10.1101/789503

**Authors:** Haisu Shi, Jinlong Tian, Chen Wu, Mo Li, Feiyu An, Rina Wu, Junhua Shao, Yan Zheng, Xue Luo, Dongbing Tao, Xu Chen, Yuzhen Pi, Chunyan Zhao, Xiqing Yue, Junrui Wu

## Abstract

Cheese lacks essential fatty acids (EFAs). Delta 12 fatty acid desaturase (*FADS12*) is a critical enzyme required for EFA biosynthesis in fermentation of the predominant strains of cheese. Previously, we identified the *FADS12* gene and characterized its function for the first time in *Geotrichum candidum*, a dominant strain used to manufacture soft cheese with white rind. In this study, we analyzed the molecular mechanism of *FADS12* function by swapping domains from *Mortierella alpina* and *G. candidum* that had, respectively, high and low oleic acid conversion rates. The results revealed three regions that are essential to this process, including regions from the end of the second transmembrane domain to the beginning of the third transmembrane domain, from the end of the third transmembrane domain to the beginning of the fourth transmembrane domain, and from the 30-amino acid from the end of the sixth transmembrane domain to the C-terminal end region. Based on our domain swapping analyses, nine pairs of amino acids including H112, S118, H156, Q161, K301, R306, E307, A309 and S323 in MaFADS12 (K123, A129, N167, M172, T302, D307, I308, E310 and D324 in GcFADS12) were identified as having a significantly effect on *FADS12* catalytic efficiency, and linoleic acid and its analogues (12,13-cyclopropenoid fatty acid) were found to inhibit the catalytic activity of *FADS12* and related recombinant enzymes. Furthermore, the molecular mechanism of *FADS12* inhibition was analyzed. The results revealed two allosteric domains, including one domain from the N-terminal region to the beginning of the first transmembrane domain and another from the 31^st^ amino acid from the end of the sixth transmembrane domain to the C terminus. Y4 and F398 amino acid residues from *MaFADS12* and eight pairs of amino acids including G56, L60, L344, G10, Q13, S24, K326 and L344 in MaFADS12 (while Y66, F70, F345, F20, Y23, Y34, F327 and F345 in GcFADS12) played a pivotal role in *FADS12* inhibition. Finally, we found that both allosteric and active sites were responsible for the catalytic activity of *FADS12* at various temperatures, pH, and times. This study offers a solid theoretical basis to develop preconditioning methods to increase the rate at which *GcFADS12* converts oleic and linoleic acids to produce higher levels of EFAs in cheese.

## Introduction

Mold cheese, fermented by multispecies microbial communities (Bertuzzi et al, 2018; Zhang et al, 2018), has been a traditional high-fat dairy product with the unsolved problem of low essential fatty acid (EFA) levels among lipids. The polyunsaturated fatty acids (PUFA) content in raw milk is relatively low in mold cheese, and its EFA level accounts for only 3.5% of total fatty acids (Hageman et al, 2019). In addition, most of the predominant fungal strains used to ferment this cheese do not have the ability to convert saturated fatty acids into many PUFAs or their conversion rate is low. Delta 12 fatty acid desaturase (*FADS12*) plays a key role in the synthesis of EFAs during fermentation. *FADS12* is a key membrane-bound desaturase that converts oleic acid [OA, 18:1^Δ9^] to linoleic acid [LA, 18:2^Δ9,12^] by introducing a double bond between carbons 12 and 13 in the carboxyl end of the substrate in the biosynthesis of EFAs (Cui et al, 2016; Lee, 2016; Rodríguez-Rodríguez, 2016). As a rate-limiting enzyme, *FADS12* plays an important role in the metabolism of PUFAs, and its activity directly affects the level and distribution of EFAs in cheese.

*FADS12* has been studied as an important desaturase in fatty acid transformation for many years, with experiments based on gene cloning, heterologous expression, and analysis of its catalytic mechanism (Brandstetter & Ruther, 2016; Kaye et al, 2015; Kazuhiro et al, 2004; Kikukawa et al, 2013; Lamers et al, 2019; Rodríguez-Rodríguez, 2016; Sakamoto et al, 2017; ShanlinYu et al, 2008; Sun et al, 2016; Watanabe et al, 2004). These studies were mostly focused on the preparation of PUFAs, the regulation of the proportion of omega-3/omega-6 fatty acids and heterologous expression from EFAs, which could not be synthesized in the human body (Wang et al, 2016; Yan et al, 2013; Zhang et al, 2013; Zhang et al, 2017). However, heterologous expression of *FADS12* in different host cells often resulted in significant changes in its activity, which has made it difficult to screen for *FADS12* with high activity in a target strain. Therefore, it is important to study differences in the mechanisms underlying differential *FADS12* activity in various species. At present, research on the catalytic mechanism of fatty acid desaturase is still in its infancy (Lee et al, 2016), except that the three-dimensional structure of *FADS9* was analyzed and found to be mushroom-shaped (Bai, 2015; Wang, 2015) and the molecular mechanisms underlying *FADS2* substrate specificity (**Shi** et al, 2015) and catalytic activity (**Shi** et al, 2018a) were analyzed in our previous study. However, studies on fatty acid desaturase in dominant strains of cheese have not yet been undertaken. Previously, our laboratory identified the *FADS12* gene in a *Geotrichum candidum* (*G. candidum*) genome sequence submitted to the GenBank database (LOCUS: CCBN010000001) by Casaregola, S. The function of this gene has been verified by the addition of substrates *in vitro*, and, although the conversion rate of the enzyme encoded by this gene has been determined, its ability to transform OA and LA substrates was found to be low (Luo et al, 2019).

In studies focusing on the catalytic mechanism of membrane-bound enzyme function, reciprocal domain swapping has commonly been used to locate regions or sites that affect activity (Aranko et al, 2013; Gammons et al, 2016; Haywood et al, 2018; Liu et al, 2012; Marcin et al, 2015; Meesapyodsuk & Qiu, 2014; Park et al, 2010; Qin et al, 2016; Sun et al, 2017; Vendome et al, 2011). We also used this method in the previous study of FADS2 to locate the domains and sites of FADS2 activity, but the conversion rates of recombinant FADS2s could not reach that of the wild-type. Therefore, we wondered whether there were any components (Bichi et al, 2012; Chen et al, 2016; Pedrono et al, 2018; Ralston et al, 2014) that hindered the synthesis of unsaturated fatty acids. R. Jeffcoat and M. R. Pollard (Jeffcoat & Pollard, 1977) pointed out in 1977 that there was an accumulation of stearic acid [8-(2-octyl-l-cyclopropenyl) octanoic acid, a C18 cyclopropenoid fatty acid] and apparent loss of OA in both tissues and lipids in cows’ milk. They showed that cyclopropenoid acid was an inhibitor of fatty acid desaturase. This is a structural analogue of other fatty acids such as 12,13-cyclopropenoid fatty acid, which might also inhibit the activity of *FADS12* and prevent the conversion of OA to LA. If so, this could help to explain the low *FADS12* activity in *G. candidum* (Luo et al, 2019) and provided a new clue to study the catalytic mechanism of fatty acid desaturase.

In the present study, key domains and amino acid sites for *FADS12* catalytic activity were located by domain swapping. In addition, the product (LA) and its analogues (12,13-cyclopropenoid fatty acid) were found to inhibit the catalytic activity of *FADS12*. Furthermore, the allosteric domains and amino acid sites that affect *FADS12* catalytic activity were identified by domain swapping. Finally, key allosteric and active domains were shown to stabilize recombinant *FADS12* at various temperatures, pH, and times.

## Materials and methods

### Domain swapping between MaFADS12 and GcFADS12

The *MaFADS12* (GeneBank accession AB020033) gene from *Mortierella alpina* was synthesized for domain swapping; *GcFADS12* (GeneBank accession MH198047) was amplified from *G. candidum* in our previous screen. *Pichia pastoris* was used for heterogeneous expression and catalytic activity determinations for *MaFADS12*, *GcFADS12*, and each chimera. Plasmid pPICZαA was used for expression of each chimera. LB agar plates and YPD medium was used as described previously (**Shi** et al, 2015). YPDS + Zeocin medium consisted of 1% yeast extract, 2% peptone, 2% glucose, 1 M sorbitol, and 100 μg/ml Zeocin. The recombinant plasmids expressing the amplified *MaFADS12* and *GcFADS12* genes were designated pPICZαA-*MaFADS12* and pPICZαA-*GcFADS12*, respectively.

To identify which structural elements play functional roles in *MaFADS12* and *GcFADS12* catabolism, both enzymes were divided into nine sections based on the transmembrane topology (hydrophilic regions and hydrophobic regions) of *FADS12* as follows: Section 1: from the N-terminal end region to the beginning of the first transmembrane domain (cytoplasmic region); Section 2: the first and the second transmembrane regions; Section 3: from the end of the second transmembrane domain to the beginning of the third transmembrane domain (cytoplasmic region); Section 4: the third transmembrane region; Section 5: the end of the third transmembrane domain to the beginning of the fourth transmembrane domain (cytoplasmic region); Section 6: the fourth transmembrane region; Section 7: the end of the fourth transmembrane domain to the beginning of the fifth transmembrane domain (cytoplasmic region); Section 8: the fifth and the sixth transmembrane regions; Section 9: the end of the sixth transmembrane domain to the C-terminal end region. In second-level domain swapping, section 5 and section 9 were divided into small sections as follows: Section 5-1: from the end of the third transmembrane domain to the 18th amino acid of section 5; Section 5-2: from the 19th amino acid of section 5 to the beginning of the fourth transmembrane domain; Section 9-1: from the end of the sixth transmembrane domain to the 30th amino acid of section 9; Section 9-2: from the 31th amino acid to the 54th (for *MaFADS12*) and 52th (for *GcFADS12*) amino acid of section 9; Section 9-3: from the 54th (for *MaFADS12*) and 52th (for *GcFADS12*) amino acid to the C-terminal end region.. All swapped recombinant genes were generated by overlap extension PCR using the primers listed in **Supplemental Table S1** and the PCR protocol described previously (**Shi** et al, 2015). The hybrid genes were ligated into the pPICZαA plasmid and transformed into competent Top10 cells.

In sections 3, 5, and 9-1, 18 mutants of the *MaFADS12* gene were constructed, with amino acids substituted with the corresponding residues in the *GcFADS12* gene (H112K, Q117G, S118A, H156N, M157L, T158Q, K159R, Q161M T298S, K301T, R306D, E307I, G308T, A309E, Q313A, L317A/C318A, V320I and S323D). Then, 18 corresponding mutants of the *GcFADS12* gene were constructed with reciprocal amino acids from the *MaFADS12* gene (K123H, G128Q, A129S, N167H, L168M, Q169T, R170K, M172Q, S299T, T302K, D307R, I308E, T309G, E310A, A314Q, A318L/A319C, I321V and D324S). Mutation primers are listed in **Supplemental Table S1**.

Each chimeric and wild-type desaturase gene was induced by 0.5 mM *cis*-OA at 28°C for 12 h under previously described conditions (**Shi** et al, 2018a). Then, the expression level of each chimeric desaturase in *Pichia pastoris* was determined by western blotting analysis as described previously (**Shi** et al, 2015); each sample loading volume was adjusted to the same level according to the western blot results to determine desaturase activity of each chimera. Chimeric desaturase activity were analyzed by gas chromatography (GC) as described previously (**Shi** et al, 2016; Shi et al, 2018b).

### Feedback inhibition of LA and its analogues

To examine *FADS12* inhibition, 0.00 mM, 0.05 mM, 0.10 mM, 0.15 mM, 0.20 mM, 0.25 mM, 0.30 mM, 0.35 mM, 0.40 mM, 0.45 mM and 0.50 mM of 12,13-cyclopropenoid fatty acid was selected as the product analogue to add to the culture medium with 0.5 mM *cis*-OA. LA, in concentrations corresponding to those of 12,13-cyclopropenoid fatty acid, was selected to verify feedback inhibition of *FADS12*. Meanwhile, the conversion rates of chimera G3, G5, G9, G3/5/9-1, chimera M3, M5, M9, and M3/5/9-1 were determined at various concentrations of 12,13-cyclopropenoid fatty acid.

### Allosteric domain swapping between chimera M3/5/9-1 and chimera G3/5/9-1

To identify which allosteric domains are functionally involved in the catalytic activities of *MaFADS12* and *GcFADS12*, the corresponding sections 1, 2, 4, 6, 7, 8, and 9-2/3 in chimeras M3/5/9-1 and G3/5/9-1 were systematically exchanged to construct recombinant swap genes that were generated by overlap extension PCR. In sections 1 and 9-2/3, 26 mutants derived from the chimeric M3/5/9-1 gene were constructed, in which amino acids were substituted with the corresponding residues in the chimeric G3/5/9-1 gene (Q34V, L35V, E37Q, R44L, E45D, C46A, H50E, F52Y, E53K, G56Y, L57V, R58K, G59S, L60F, G334D, V336I, H337E, A341L, L344F, F345V, Q347R, G10F, Q13Y, S24Y, K326F and L344F). Then, 28 corresponding mutants derived from the chimeric G3/5/9-1 gene were constructed, in which amino acids were substituted with the corresponding residues in the chimeric M3/5/9-1 gene (V44Q, V45L, Q47E, L54R, D55E, A56C, E60H, Y62F, K63E, Y66G, V67L, K68R, S69G, F70L, D335G, I337V, E338H, L342A, F345L, V346F, R348Q, Y4A, F20G, Y23Q, Y34S, F327K, F345L and F398A). Mutation primers are listed in **Supplemental Table S1**.

### Determination of chimeric FADS12 properties

Chimera M3/5/9-1, chimera G3/5/9-1, chimera Al M1/9-2/3, chimera Al G1/9-2/3, *MaFADS12*, and *GcFADS12* were selected to determine catalytic properties, including induction temperatures (18°C, 23°C, 28°C, 33°C, and 38°C at pH 7.0 with a 12-h induction), pH (6.0, 6.5, 7.0, 7.5, and 8.0 at 28°C with a 12 h induction), and induction times (4h, 8 h, 12 h, 16 h, 20 h, and 24 h at pH 7.0 and 28°C).

### Molecular docking of fatty acids on the binding site of FADS12

The Discovery Studio 4.3 program (Accelrys Inc., San Diego, CA, USA) was used to optimize *FADS12* by adding hydrogen atoms and removing water molecules, and the Molegro. Docker 4.0 program (Molegro ApS, Aarhus, Denmark) was used to prepare the *FADS12* structure for docking calculations.

## Results

### Locating key domains and amino acid sites associated with FADS12 catalytic activity

Based on a multiple sequence alignment of the *FADS12*s and their C18:1 catalysis efficiencies **(Fig. S1)**, *MaFADS12* with highest catalytic activity and *GcFADS12* with lowest catalytic activity were selected for domain swapping. The results of domain swapping showed that the catalytic efficiencies of chimera M3 (*MaFADS12* aa111–120 replaced by *GcFADS12* aa122–131), chimera M5 (*MaFADS12* aa150–190 replaced by *GcFADS12* aa161–197), and chimera M9 (*MaFADS12* aa295–400 replaced by *GcFADS12* aa296–412) were 12.6 ± 0.7%, 15.9 ± 0.6%, and 11.3 ± 1.6% respectively, much lower than those of wild-type *MaFADS12* (87.6 ± 0.6%). Conversely, whereas *GcFADS12* exhibited a catalytic efficiency of 20.4 ± 1.0%, those of chimera G3 (*GcFADS12* aa122–131 replaced by *MaFADS12* aa111–120), chimera G5 (*GcFADS12* aa161–197 replaced by *MaFADS12* aa150–190), and chimera G9 (*GcFADS12* aa296–412 replaced by *MaFADS12* aa295–400) were increased to 44.3 ± 0.5%, 28.2 ± 0.5%, and 46.5 ± 4.0%, respectively (Figure 1). Furthermore, section 9 was divided into three ‘second-level’ sections to identify key functional areas that were then systematically exchanged between *MaFADS12* and *GcFADS12*. The results of an analysis of the second-level chimeras showed that the catalytic efficiency of chimera M9-1 (aa295–325 in *MaFADS12* replaced by aa296–326 in *GcFADS12*) was decreased to 5.6 ± 1.9%, compared to that exhibited by *MaFADS12* (Figure 1). Likewise, replacement of aa296–326 (section 9-1) of *GcFADS12* with aa295–325 of *MaFADS12* resulted in significantly increased (36.6 ± 1.0%) catalytic efficiency (Figure 1). Replacement of the other two second-level sections (sections 9-2 and 9-3) had no significant impact on the catalytic efficiency of *FADS12*. Key functional areas, namely sections 3, 5, and 9-1, were exchanged simultaneously between *MaFADS12* and *GcFADS12*, and our results showed that the catalytic efficiency of chimera M3/5/9-1 decreased to 5.6 ± 0.6%, whereas that of chimera G3/5/9-1 reached 56.3 ± 2.3%, exhibiting either lower or higher efficiencies than those of the other corresponding section (i.e., the catalytic efficiencies of chimeras M3, 5, and 9-1 in *MaFADS12* were lower than those of chimeras G3, 5, and 9-1 in *GcFADS12*).

**Figure 1.**
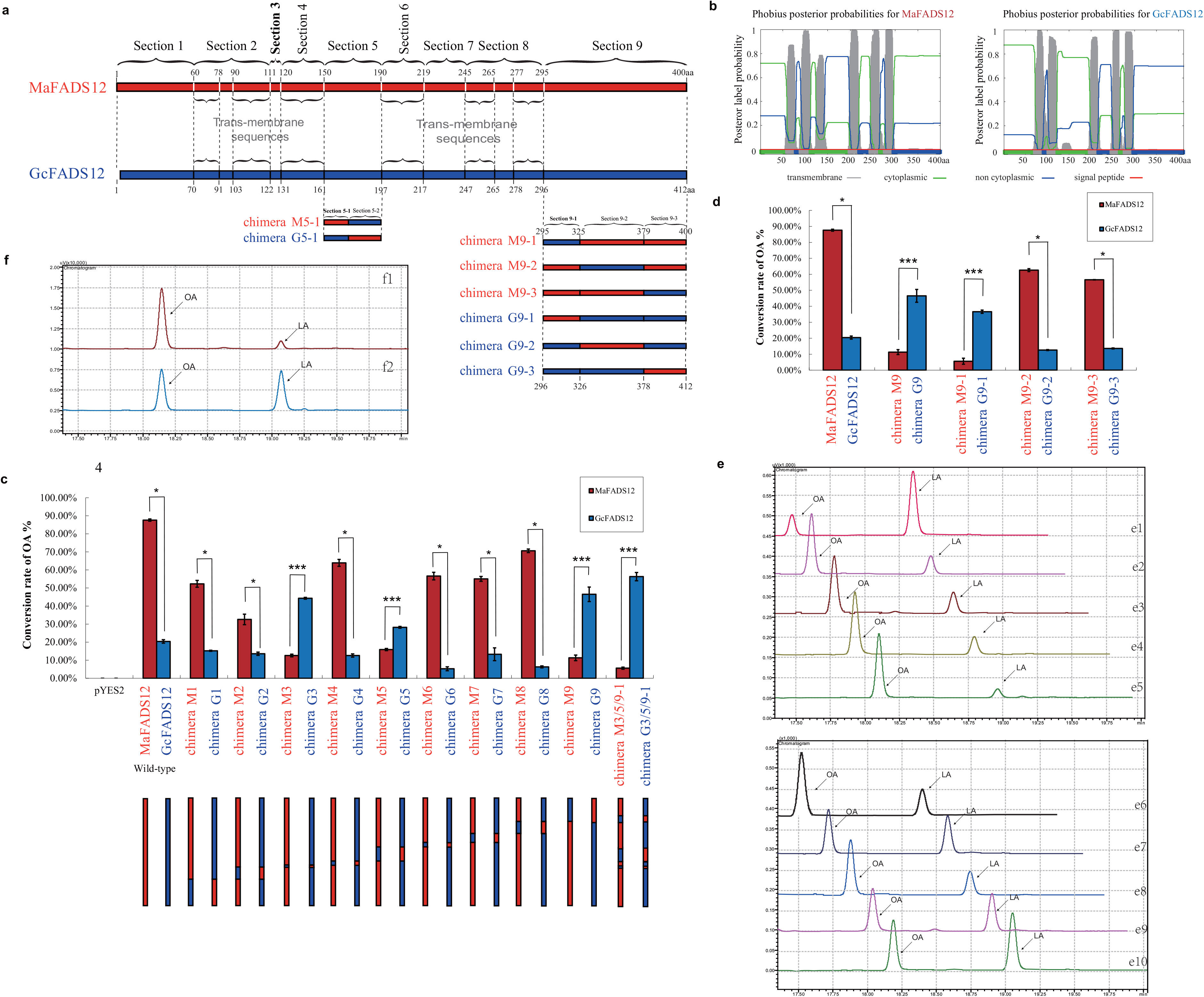
(a): Skeleton map of the amino acid sequences of MaFADS12 (red box) and GcFADS12 (blue box), which were divided into 9 sections showed with dotted lines. The braces indicate their trans-membrane sequences. Section 5 was divided into 2 sections and section 9 was divided into 3 sections with dotted lines. (b): phobius posterior probalilities for MaFADS12 and GcFADS12. (c): Relative substrate conversion efficiency of each FADS12 chimera expressed in *Pichia.pastoris*, determined by adding LA substrate. Map of each FADS12 chimera constructed by reciprocal section swapping was exhibited below each FADS12 chimera name. Substrate conversion efficiency of each chimera was classified into two groups: *, conversion rate of chimera M*X* > chimera G*X*; ***, conversion rate of chimera M*X* < chimera G*X*. (d): Relative substrate conversion efficiency of each second-level chimera in section 9 expressed in *Pichia.pastoris* determined by adding LA substrate. (e): Gas chromatogram of fatty acids for the conversion rate of MaFADS12 (e1), chimera M3 (e2), chimera M5 (e3), chimera M9 (e4), chimera M3/5/9-1 (e5), GcFADS12 (e6), chimera G3 (e7), chimera G5 (e8), chimera G9 (e9) and chimera G3/5/9-1 (e10). (f): Gas chromatogram of fatty acids for the conversion rate of chimera M9-1 (f1) and chimera G9-1 (f2).

To further identify key amino acid sites associated with *FADS12* catalytic activity, a series of constructs with mutations in sections 3, 5, and 9-1 were generated (Figure 2). Conversion efficiencies of these constructs were shown to be greatly reduced in single mutants H112K, S118A, H156N, and Q161M to a rate of 23.6 ± 1.6 %, 25.6 ± 2.5 %, 20.9 ± 3.9 %, and 22.6 ± 2.1 %, respectively, for *MaFADS12* with mutant sections 3 and 5 (Figure 2). Conversely, catalytic efficiencies were significantly enhanced in single mutants K123H, A129S, N167H, and M172Q (35.2 ± 1.3 %, 43.0 ± 2.1 %, 32.6 ± 3.5 %, and 36.3 ± 2.7 %, respectively, for *GcFADS12* with mutant sections 3 and 5) (Figure 2). The conversion efficiencies of the section 9-1 *MaFADS12* mutants K301T, R306D, E307I, A309E, and S323D and the corresponding *GcFADS12* mutants T302K, D307R, I308E, E310A, and D324S showed significant differences from those of wild-type *FADS12* (Figure 2). Targeted mutagenesis of other points within sections 3, 5 and 9-1 did not induce any major changes in *FADS12* catalytic activity.

**Figure 2.**
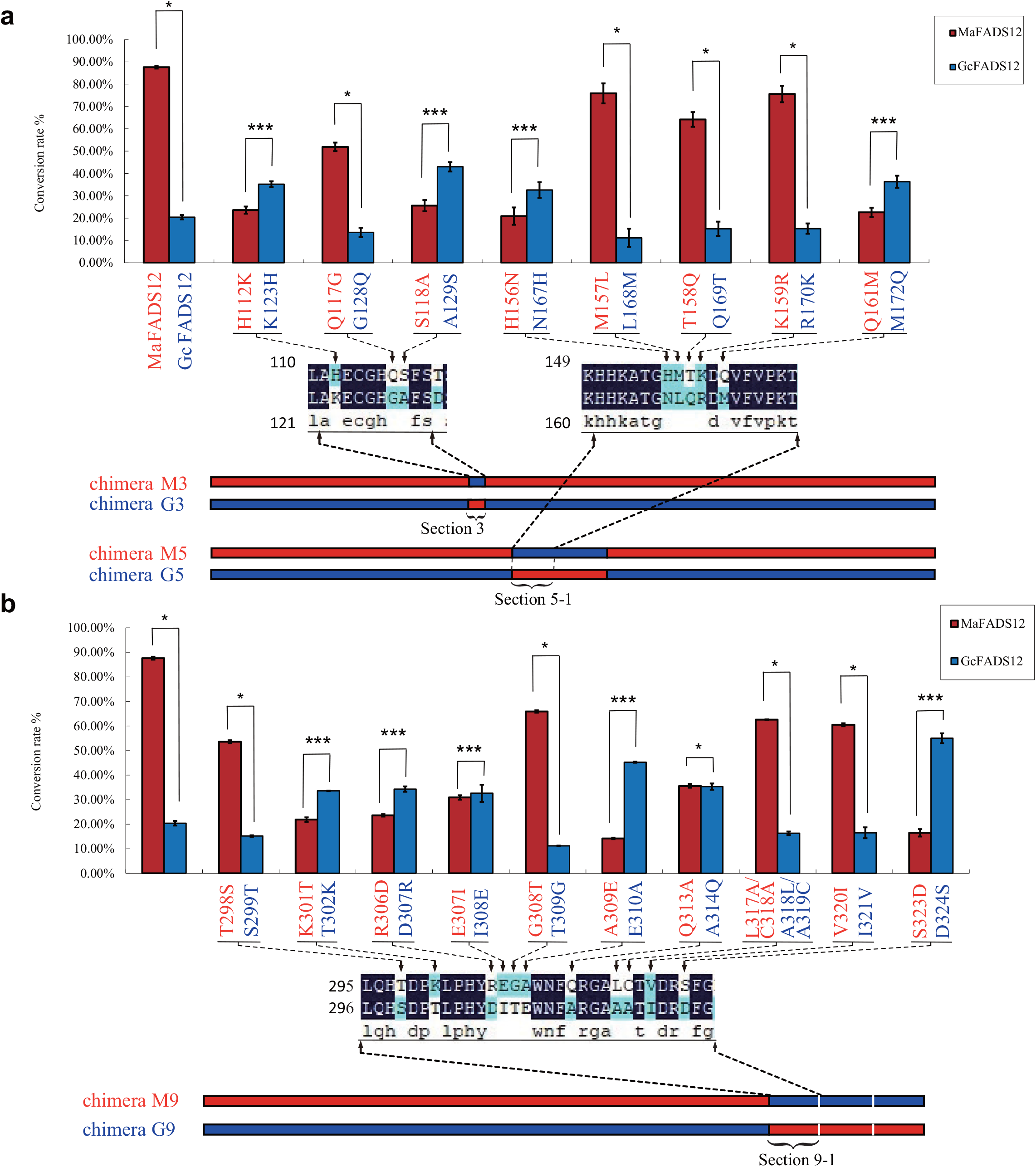
Relative substrate conversion efficiencies of each mutant from section 3, 5-1 and 9-1. The mutants (red words under red column) of H112K, Q117G, S118A, H156N, M157L, T158Q, K159R, Q161M T298S, K301T, R306D, E307I, G308T, A309E, Q313A, L317A/C318A, V320I and S323D from MaFADS12 were corresponding to the mutants (blue words under blue column) of K123H, G128Q, A129S, N167H, L168M, Q169T, R170K, M172Q, S299T, T302K, D307R, I308E, T309G, E310A, A314Q, A318L/A319C, I321V and D324S from GcFADS12, respectively. Substrate conversion efficiency of each mutant was classified into two groups: *, conversion rate of mutant from MaFADS12 > mutant from GcFADS12; ***, conversion rate of mutant from MaFADS12 < mutant from GcFADS12. (a): each mutant from section 3 and 5-1; (b): each mutant from section 9-1.

### Inhibition of FADS12 catalytic activity by 12,13-cyclopropenoid fatty acid

To explain why the increase in activity of chimeras with key regions and sites described above was not significant, we considered that there may be reaction components that inhibited activity of the enzyme. Therefore, we explored feedback inhibition. By adding LA or 12,13-cyclopropenoid fatty acid (an LA analogue) as suspected inhibitors of enzyme activity or a control (with neither fatty acid), we showed that both LA and 12,13-cyclopropenoid fatty acid inhibited *FADS12* activity (Figure 3a, b, c). This result was confirmed by the effect of LA or 12,13-cyclopropenoid fatty acid on *GcFADS12* activity at a low concentration of LA or 12,13-cyclopropenoid fatty acid with substrate in the medium. *MaFADS12* activity did not significantly change when OA and 12,13-cyclopropenoid fatty acid or OA and LA were added, whereas *GcFADS12* activity decreased with an increase in 12,13-cyclopropenoid fatty acid or LA. When the concentration of 12,13-cyclopropenoid fatty acid or LA reached 0.5 mM, *GcFADS12* activity decreased from 20.4 % to less than 3 %, which confirmed that these compounds strongly inhibit *GcFADS12*.

**Figure 3.**
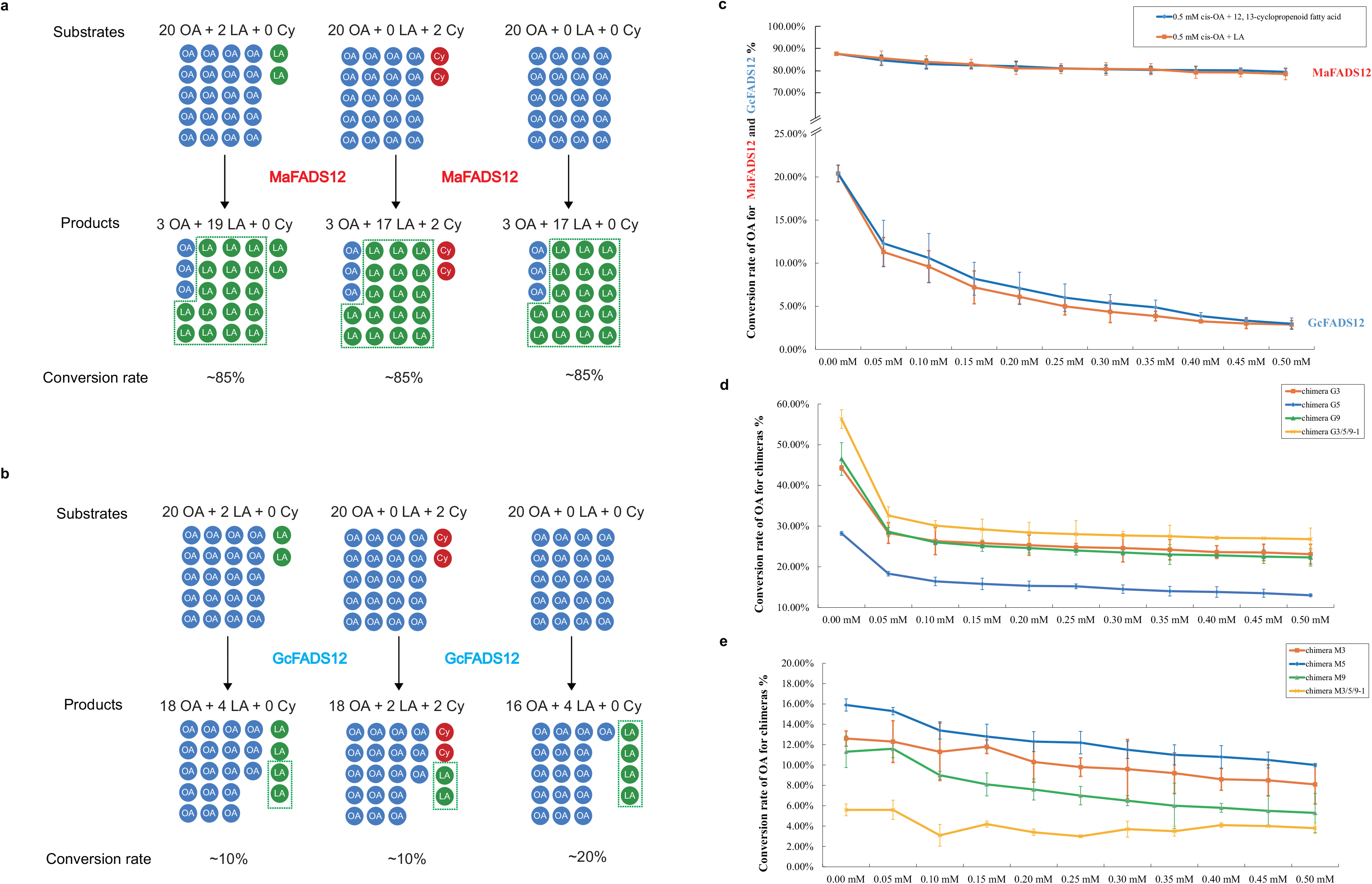
Feedback inhibition of LA and its analogues for MaFADS12 and GcFADS12. (a): Map of catalytic reaction of MaFADS12 by LA addition, 12, 13-cyclopropenoid fatty acid addition and control. (b): Map of catalytic reaction of GcFADS12 by LA addition, 12, 13-cyclopropenoid fatty acid addition and control. (c): Relative substrate conversion efficiency of MaFADS12 and GcFADS12 by addition of 0.5 mM *cis*-OA/various concentrations of 12, 13-cyclopropenoid fatty acid and 0.5 mM *cis*-OA/various concentrations of LA, respectively. (d): Relative substrate conversion efficiency of chimera G3, G5, G9 and G3/5/9-1 by addition of 0.5 mM *cis*-OA/various concentrations of 12, 13-cyclopropenoid fatty acid. (e): Relative substrate conversion efficiency of chimera M3, M5, M9 and M3/5/9-1 by addition of 0.5 mM *cis*-OA/various concentrations of 12, 13-cyclopropenoid fatty acid. Substrate conversion efficiency with LA addition had removed the level of added LA.

Moreover, the effects of 12,13-cyclopropenoid fatty acid on the activity of chimeras G3, G5, G9, G3/5/9-1, M3, M5, M9, and M3/5/9-1 (shown by domain swapping) were analyzed. Our results showed that activities of four *GcFADS12* mutant enzymes, namely chimeras G3, G5, G9, and G3/5/9-1) decreased to half that of *GcFADS12* at a concentration as low as 0.05 mM of 12,13-cyclopropenoid fatty acid. With an increase in concentration, the activity of these chimeras showed a slight decline (Figure 3d). This confirmed that 12,13-cyclopropenoid fatty acid affected the activities of these four chimeras, and that the concentration of LA or 12,13-cyclopropenoid fatty acid was not the real reason for the change in enzyme activities. However, the activities of the four chimeras from *MaFADS12* (chimeras M3, M5, M9, and M3/5/9-1) remained unchanged in the presence of 12,13-cyclopropenoid fatty acid at various concentrations (Figure 3e). These results indicated that there may be “sensitive” regions for 12,13-cyclopropenoid fatty acid or LA in the structure of *GcFADS12* (“sensitive” regions, meaning regions that were changed by the product and that affected binding between *GcFADS12* and the substrate), and that 12,13-cyclopropenoid fatty acid does not change the corresponding binding regions in *MaFADS12*.

### Allosteric domains and amino acid sites also affect FADS12 catalytic activity

To explain the effects of 12,13-cyclopropenoid fatty acid and LA on *GcFADS12* activity, we speculated that the product changed the structure outside the active domains of *GcFADS12*, whereas the product did not change or changed only to a small extent the corresponding region in *MaFADS12*. To locate allosteric domains, we swapped domains other than those with localized active regions (i.e., chimeras 1, 2, 4, 6, 7, 8, and 9-2/3). The results showed that the rates of conversion to OA by chimeras Al M1 and Al G1 derived from section 1 of *MaFADS12* and *GcFADS12* were 0.5 ± 0.2% (10 times lower than that of chimera M3/5/9-1) and 75.2 ± 0.3% (about 20% higher than that of chimera G3/5/9-1), respectively (Figure 4a). In addition, the rates of conversion to OA chimeras Al M9-2/3 and Al G9-2/3 derived from section 9-2/3 of *MaFADS12* and *GcFADS12* were 0.3 ± 0.1% (22 times lower than that of chimera M3/5/9-1) and 73.3 ± 3.5% (about 18% higher than that of chimera G3/5/9-1) (Figure 4a). This result suggested that there were two regions in *MaFADS12*, section 1 and section 9-2/3 that housed allosteric domains. Moreover, the conversion rates of chimeras Al M1/9-2/3 and Al G1/9-2/3, which included the section 1 and 9-2/3 domains of *MaFADS12* and *GcFADS12*, respectively, were 0.1 ± 0.0% and 80.7 ± 5.6%, which were higher than those observed with single domain swapping (Figure 4a). This confirmed that these two regions were the root cause of feedback inhibition.

**Figure 4.**
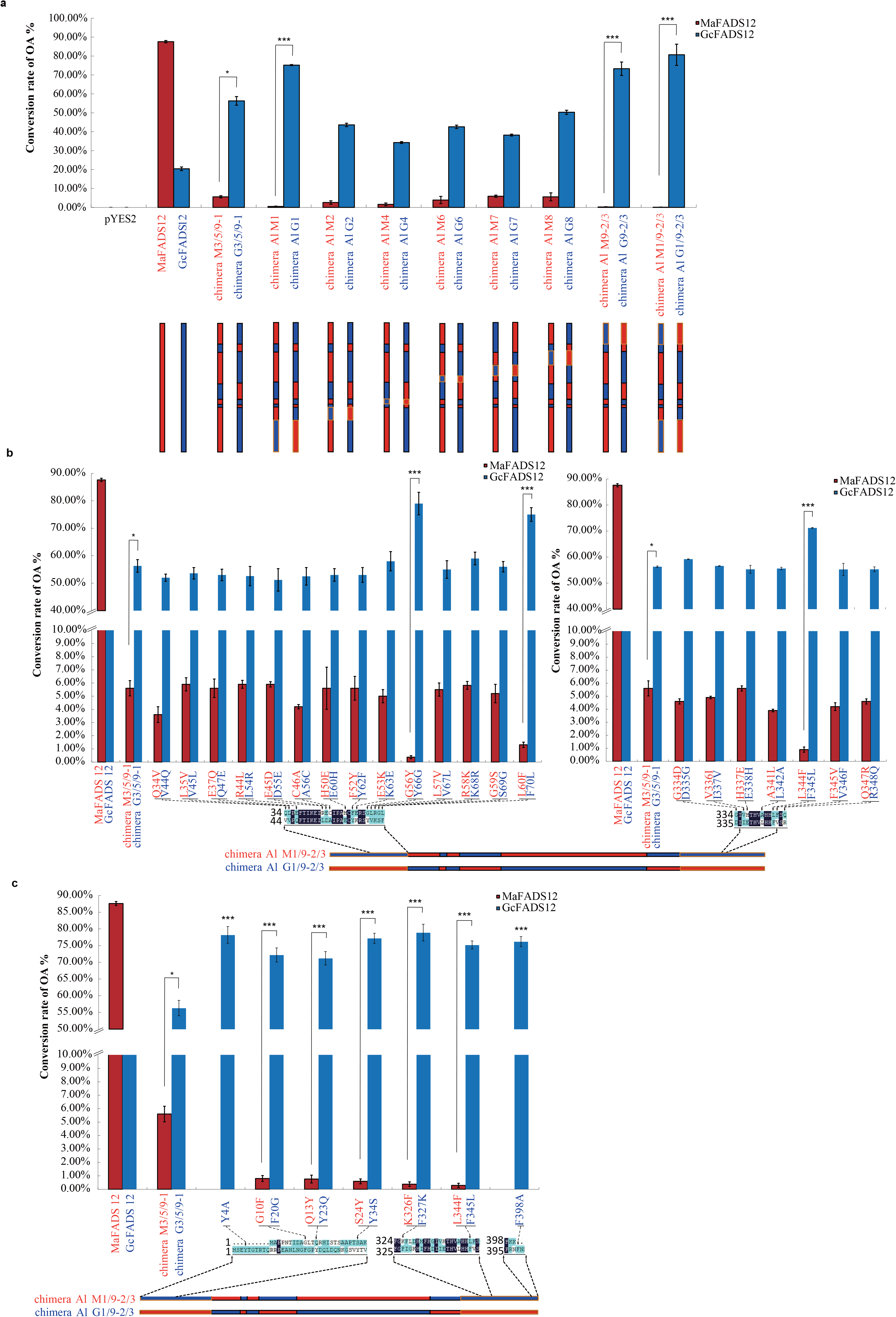
(a): Relative substrate conversion efficiencies of each FADS12 chimera constructed from chimera M3/5/9-1 and chimera G3/5/9-1 expressed in *Pichia.pastoris*, determined by adding LA substrate. Map of each chimera constructed by reciprocal section swapping was exhibited below each FADS12 chimera name. Substrate conversion efficiency of each chimera was classified into two groups: *, conversion rate of chimera M3/5/9-1 and chimera G3/5/9-1; ***, conversion rate of chimera Al M*X* < chimera M3/5/9-1 and chimera Al G*X* > chimera G3/5/9-1 chimera G*X*. (b) and (c): Relative substrate conversion efficiency of each mutant from chimera M3/5/9-1 and chimera G3/5/9-1, determined by adding LA substrate. The mutants (red words under red column) of Q34V, L35V, E37Q, R44L, E45D, C46A, H50E, F52Y, E53K, G56Y, L57V, R58K, G59S, L60F, G334D, V336I, H337E, A341L, L344F, F345V, Q347R, G10F, Q13Y, S24Y, K326F and L344F from chimera M3/5/9-1 were corresponding to the mutants (blue words under blue column) of V44Q, V45L, Q47E, L54R, D55E, A56C, E60H, Y62F, K63E, Y66G, V67L, K68R, S69G, F70L, D335G, I337V, E338H, L342A, F345L, V346F, R348Q, Y4A, F20G, Y23Q, Y34S, F327K, F345L and F398A from chimera G3/5/9-1, respectively. Substrate conversion efficiency of each mutant was classified into two groups: *, conversion rate of chimera M3/5/9-1 and chimera G3/5/9-1; ***, conversion rate of mutants from chimera M3/5/9-1 < chimera M3/5/9-1 and mutants from chimera G3/5/9-1 > chimera G3/5/9-1.

To further define key sites of feedback inhibition in the two regions, we first selected sites near conserved regions in the allosteric domain for site-directed mutagenesis swaps in *MaFADS12* and *GcFADS12*. Our results showed that the conversion rates of mutants G56Y and L60F with section 1 from *MaFADS12* were reduced to 0.4 ± 0.1% and 1.3 ± 0.2%, respectively, whereas the conversion rates of the corresponding mutants Y66G and F70L with section 1 from *GcFADS12* increased to 79.0 ± 4.1% and 75.0 ± 2.5%, respectively (Figure 4b). In addition, the conversion rates of mutant L344F with section 9-2/3 from *MaFADS12* and the corresponding F345L mutant with section 9-2/3 from *GcFADS12* significantly changed, reaching 0.9 ± 0.2% and 71.2 ± 0.5%, respectively (Figure 4b).

Based on the results of this site-directed mutagenesis swap, tyrosine and phenylalanine, aromatic amino acids with a benzene ring structure, were implicated in *FADS12* catabolism. We inferred that amino acids with large steric hindrances such as benzene rings had a greater impact on product feedback inhibition. Subsequently, other aromatic amino acids in sections 1 and 9-2/3 of *GcFADS12* were selected for exchange with corresponding sites in *MaFADS12*. Our results showed that the conversion rates of all *MaFADS12* mutants, including G10F, Q13Y, S24Y, K326F, and L344F, were reduced by an order of magnitude compared with those of chimera M3/5/9-1 (0.8 ± 0.2%, 0.8 ± 0.3%, 0.6 ± 0.2%, 0.4 ± 0.2%, and 0.3 ± 0.2%, respectively), whereas the conversion rates of Y4A, F398A, and corresponding mutants from *GcFADS12*, namely F20G, Y23Q, Y34S, F327K, and F345L, increased by about 20% compared with those of chimera G3/5/9-1, reaching 78.2 ± 2.5%, 72.2 ± 2.1%, 71.2 ± 2.0%, 77.2 ± 1.5%, 78.9 ± 2.5%, 75.2 ± 1.2%, and 76.2 ± 1.5%, respectively (Figure 4c). This result indicated that the aromatic amino acids in sections 1 and 9-2/3 of *GcFADS12* caused feedback inhibition that affected the catalytic activity of *FADS12*.

### Key allosteric and active domains with FADS12-stabilizing properties

Based on these results, we determined that there are both active and allosteric regions in *MaFADS12* that are associated with high catalytic activity, and that product feedback inhibition was weak with this enzyme because there were no aromatic amino acids in the allosteric region so there was little steric hindrance. To determine whether this conclusion was valid under different conditions, the conversion rates of *MaFADS12*, *GcFADS12*, and their recombinant enzymes at various induction temperatures, pH, and times were determined. Our results showed that the conversion rates of *MaFADS12*, chimera M3/5/9-1, and chimera Al G1/9-2/3 did not change significantly at different induction temperatures, whereas those of *GcFADS12*, chimera G3/5/9-1, and chimera Al M1/9-2/3 fluctuated with temperature (Figure 5a). The same conclusion can be drawn about substrate conversion efficiencies at various pH and induction times (Figure 5b, c). From this result, it is apparent that the conversion efficiencies of *FADS12*s containing allosteric regions (i.e., *MaFADS12*, chimera M3/5/9-1, and chimera Al G1/9-2/3) were not influenced by induction temperature, pH, or time, but for *FADS12*s without allosteric regions, even if they contained active regions (i.e., *GcFADS12*, chimera G3/5/9-1, and chimera Al M1/9-2/3), their conversion rates were unstable with a change in induction temperature, pH, or time (Figure 5). Based on these results, we knew that two key allosteric and active domains stabilize *FADS12* properties.

**Figure 5.**
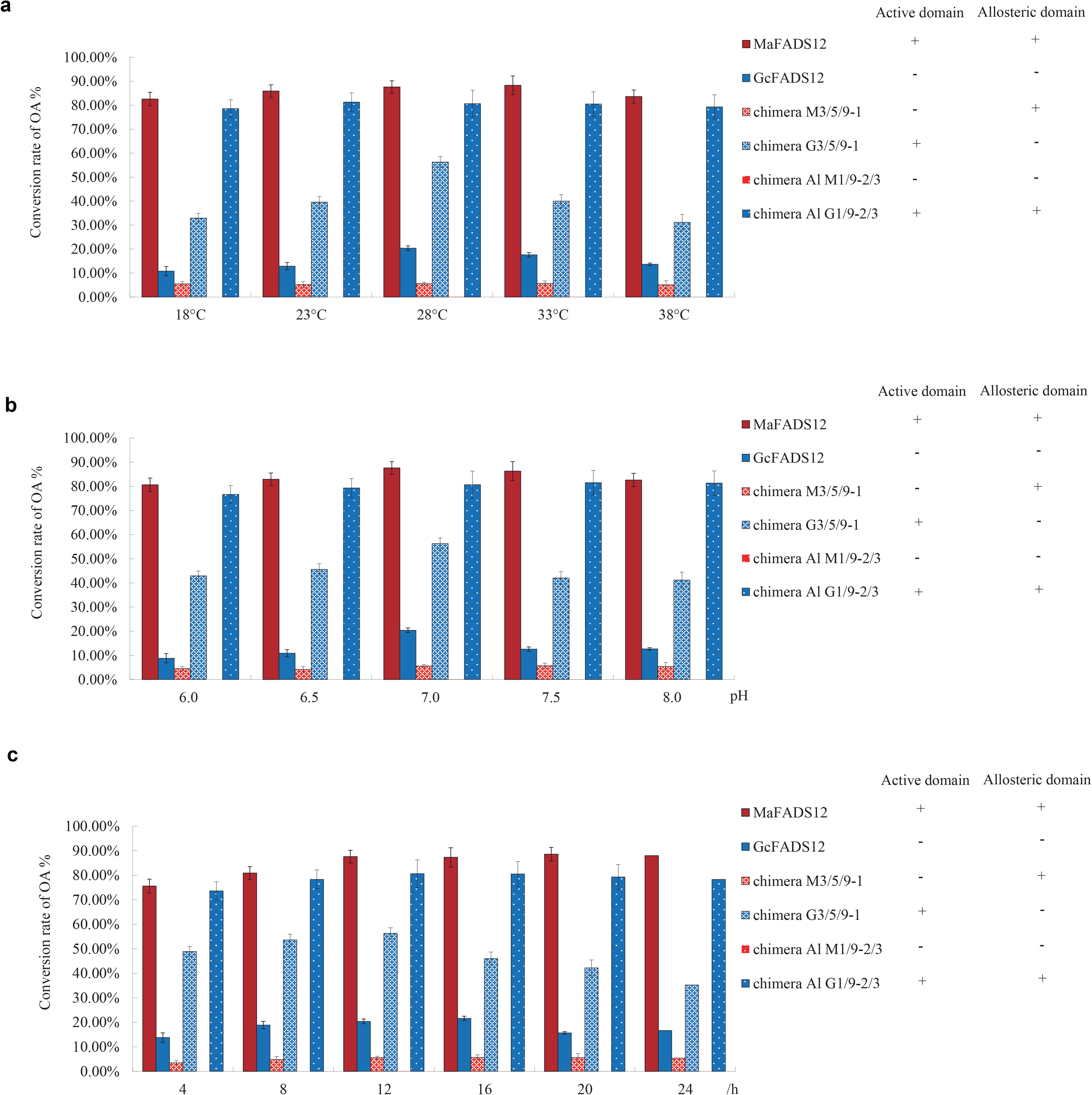
Relative substrate conversion efficiencies of chimera M3/5/9-1, chimera G3/5/9-1, chimera Al M1/9-2/3, chimera Al G1/9-2/3, MaFADS12 and GcFADS12 at various induced temperature (a), pH (b) and induced time (c). Active domain and Allosteric domain in chimera M3/5/9-1, chimera G3/5/9-1, chimera Al M1/9-2/3, chimera Al G1/9-2/3, MaFADS12 and GcFADS12 indicated with “+”, none with “-”.

## Discussion

The present study indicates that *FADS12* cytoplasmic regions extending from the end of the second transmembrane domain to the beginning of the third transmembrane domain (section 3), the end of the third transmembrane domain to the 18th amino acid of section 5 (section 5-1), and the end of the sixth transmembrane domain to the 30th amino acid of section 9 (section 9-1) critically mediates *FADS12* catalytic activity (Figure 1). In these three small regions, 10 sites (His112, Ser118, His156, Gln161, Lys 301, Arg306, Glu307, Ala 309, Gln313 and Ser323) in *MaFADS12* and 10 sites (Lys123, Ala129, Asn167, Met172, Thr302, Asp307, Ile308, Glu310, Ala314 and Asp324) in *GcFADS12* were identified as key sites affecting *FADS12* catalytic activity. From the results of molecular docking experiments with these sites and the substrate, we learned that all key sites were close to the substrate, and, in the enzyme pocket, these sites associated with the substrates by van der Waals and conventional hydrogen bonds to form a stable structure (Figure 6a). The results of molecular docking experiments were in full agreement with the observed *FADS12* conversion rates of *FADS12* (Figure 6b).

**Figure 6.**
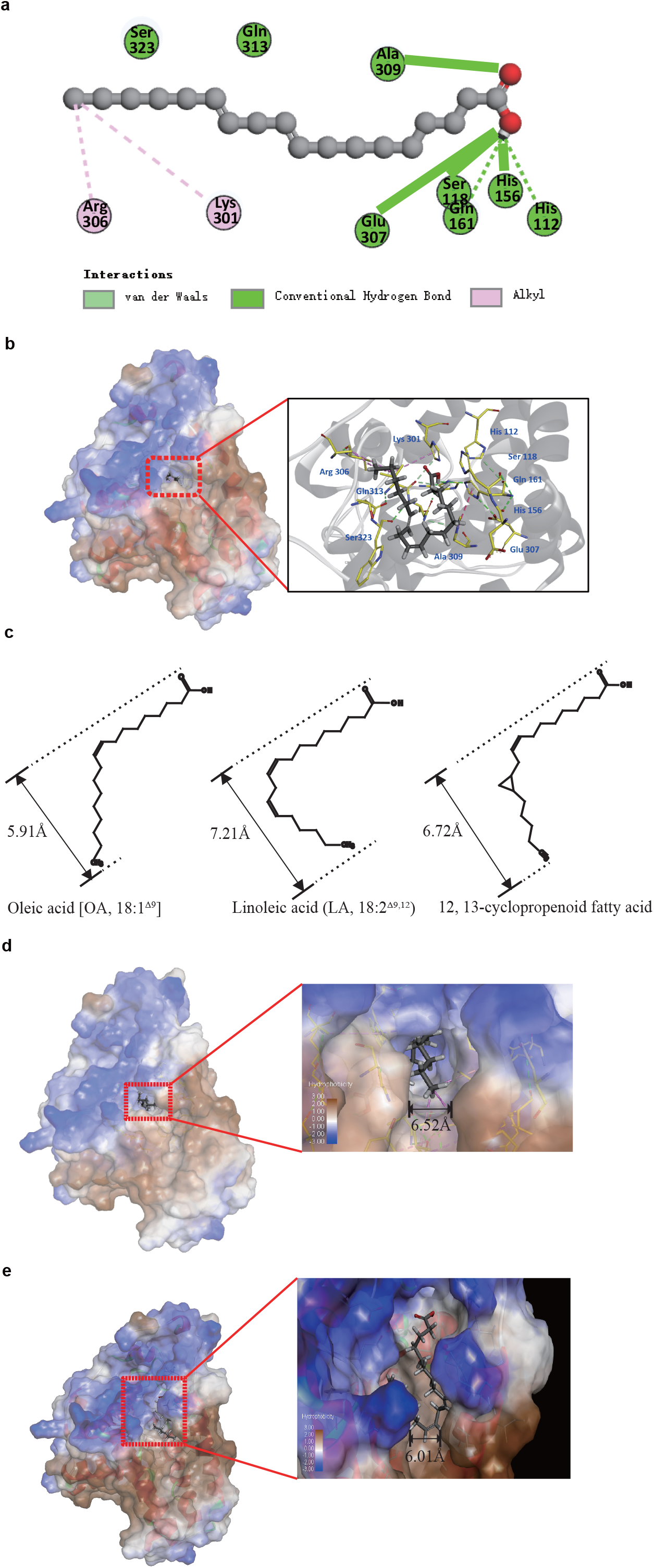
(a): H-bonds and hydrophobic interactions between LA and MaFADS12 in a 2D docking model. (b): LA interacted with the amino acid residues located within the active site of MaFADS12 pocket in 3D docking model. (c): The molecular structure of OA, LA and 12,13-cyclopropenoid fatty acid with diameters perpendicular to the direction of the substrate entering the pocket of FADS12. (d): Predicted binding mode of LA docked into key amino acids of MaFADS12. The binging location of MaFADS12 interacting with LA was shown as molecular surface structures; (e): Predicted binding mode of LA docked into key amino acids mutated to tyrosine and phenylalanine in MaFADS12 pocket.

The conversion rate of chimera G3/5/9-1, constructed by superimposition of three small active domains (sections 3, 5-1, and 9-1), reached 56.3 ± 2.3%, which was 30% lower than that of wild-type *MaFADS12*. We speculated that this was due to feedback inhibition by the product. To exclude the effect of products on the conversion rate, 12,13-cyclopropenoid fatty acid was used as a product analogue; this confirmed that 12,13-cyclopropenoid fatty acid inhibited *GcFADS12* activity, but had no significant effect on *MaFADS12* activity. There are likely allosteric regions in *MaFADS12* that are less sensitive to the product, whereas this allosteric region in *GcFADS12* is highly sensitive to it. In addition, the minimal steric hindrance of OA and maximal steric hindrance of LA and 12,13-cyclopropenoid fatty acid in various twisted structures were analyzed, and diameters at which the said hindrance may interfere were 5.91Å, 7.21Å, and 6.72Å respectively (Figure 6c). By comparing the pocket size of the active region of *FADS12* with the steric hindrance distance of the product or its analogues, we inferred that it was difficult for LA to be released from the pocket due to steric hindrance after the catalysis of the OA substrate and feedback inhibition by the product (Figure 6d).

Allosteric regions were again found to be located in chimeras M3/5/9-1 and G3/5/9-1 after domain swapping. The results of this study showed that the activities of chimeras Al G1 and Al G9-2/3, both of which were derived from chimera G3/5/9-1 by exchanging the first and last regions, respectively, were close to those of wild-type *MaFADS12*; and the activity of chimera Al G1/9-2/3, which was derived by exchanging the first and last regions, was similar to that of wild-type *MaFADS12* (Figure 4a). Based on the simulated structure of *MaFADS12*, these two segments can be considered “flexible regions,” which change during the catalytic reaction to release the product (Figure 6d). In these two regions, once an amino acid with a benzene ring structure was mutated in *GcFADS12*, its activity significantly increased, whereas activity in the corresponding site of *MaFADS12* significantly decreased (Figure 4b, c). From the docking results, we inferred that macromolecule groups such as a benzene ring at the first and last regions in *FADS12* hindered product release (Figure 6e).

The substrate conversion rate is determined by both active and allosteric regions. With changes in the induction temperature, pH, or time, the substrate conversion rates of recombinant *FADS12* containing a "flexible region" (i.e., *MaFADS12*, chimera M3/5/9-1, and chimera Al G1/9-2/3) showed few significant changes, indicating that the flexible region plays a role in product release, even when the overall three-dimensional structure of the enzyme changed with external conditions. However, the conversion rate of chimera G3/5/9-1 with an active region but no allosteric region was not stable under different conditions, suggesting that this flexible region is the key domain to maintaining *FADS12* stability (Figure 5). Here, the three-dimensional structure of the two regions could be analyzed by X-ray diffraction in the future, which will help elucidate how the two regions combine with the substrate.

## Conclusions

The present study elucidated the molecular mechanism underlying the catalytic activity of *FADS12* from *M. alpina* and *G. candidum*. A catalytic product of *FADS12* was shown to be an inhibitor of *FADS12* catabolism; also, for the first time, a product analogue as found to inhibit this enzyme. The molecular mechanism underlying *FADS12* inhibitory activity was analyzed based on the presence of allosteric domains and amino acid sites affecting *FADS12* catalytic activity. These studies indicated that both allosteric and active domains can stabilize *FADS12* in various induction conditions. This study provides a theoretical basis for improving essential fatty acid contents in dominant strains in fermented cheese.

## Acknowledgments

This study was supported by the National Natural Science Foundation of China (No. 31801567, 31871831 and 31470538), General Project of the Educational Department of Liaoning Province of China (LSNQN201724), the PhD Start-up Fund of Shenyang Agricultural University and Tianzhushan Excellent Talents program of Shenyang Agricultural University.

---

**Fig. S1** Multiple Sequence alignment of deduced amino acids for GcFADS12 with FADS12s from other species, including *Mortierella alpina*, MaFADS12, *Mucor circinelloides*, McFADS12, *Cannabis sativa*, CsFADS12, *Arachis hypogaea*, AhFADS12, *Santalum acuminatum*, SaFADS12, *Spinacia oleracea*, SoFADS12, *Camelina sativa*, CmFADS12, *Fusarium verticillioides*, FvFADS12, *Candida parapsilosis*, CpFADS12, *Exocarpos cupressiformis*, EcFADS12 using DNAMAN software. Black line indicate domain boundaries for reciprocal section swapping. CA, catalytic activity.

**Fig. S2** Gas chromatogram of fatty acids in *Pichia.pastoris* carrying MaFADS12 (a), GcFADS12 (b) genes and each substitution including H112K (c), S118A (e), H156N (g), Q161M (i), K301T (k), R306D (m), E307I (o), A309E (q), Q313A (s) and S323D (u) from MaFADS12 and K123H (d), A129S (f), N167H (h), M172Q (j), T302K (l), D307R (n), I308E (p), E310A (r), A314Q (t) and D324S (v) from GcFADS12 with 0.5 mM *cis*-OA addition at 28°C for 12 h induction.

**Fig. S3** Gas chromatogram of fatty acids in *Pichia.pastoris* carrying MaFADS12 (a), GcFADS12 (b) genes and each substitution including G56Y (c), L60F (e), L344F (g), G10F (i), Q13Y (k), S24Y (m), K326F (o) and L344F (q) from MaFADS12 and Y66G (d), F70L (f), F345L (h), F20G (j), Y23Q (l), Y34S (n), F327K (p), F345L (r), Y4A (s) and F398A (t) from GcFADS12 with 0.5 mM *cis*-OA addition at 28°C for 12 h induction.

## References

Aranko AS, Oeemig JS, Kajander T, Iwaï H (2013) Intermolecular domain swapping induces intein-mediated protein alternative splicing. Nature Chemical Biology 9: 616–622

Bai Y, McCoy, Jason G., Levin, Elena J., Sobrado, Pablo, Rajashankar, Kanagalaghatta R., Fox, Brian G., Zhou, Ming (2015) X-ray structure of a mammalian stearoyl-CoA desaturase. Nature 524: 252–256

Bertuzzi AS, McSweeney PLH, Rea MC, Kilcawley KN (2018) Detection of Volatile Compounds of Cheese and Their Contribution to the Flavor Profile of Surface-Ripened Cheese. Compr Rev Food Sci Food Saf 17: 371–390

Bichi E, Toral PG, Hervas G, Frutos P, Gomez-Cortes P, Juarez M, de la Fuente MA (2012) Inhibition of 9-desaturase activity with sterculic acid: effect on the endogenous synthesis of cis-9 18:1 and cis-9, trans-11 18:2 in dairy sheep. J Dairy Sci 95: 5242–5252

Brandstetter B, Ruther J (2016) An insect with a delta-12 desaturase, the jewel wasp Nasonia vitripennis, benefits from nutritional supply with linoleic acid. Die Naturwissenschaften 103: 40

Chen F, Di H, Wang Y, Cao Q, Xu B, Zhang X, Yang N, Liu G, Yang CG, Xu Y (2016) Small-molecule targeting of a diapophytoene desaturase inhibits S. aureus virulence. Nature Chemical Biology 12: 174–179

Cui J, He S, Ji X, Lin L, Wei Y, Zhang Q (2016) Identification and characterization of a novel bifunctional Δ12/Δ15-fatty acid desaturase gene from *Rhodosporidium kratochvilovae*. Biotechnology letters 38: 1155–1164

Gammons MV, Renko M, Johnson CM, Rutherford TJ, Bienz M (2016) Wnt Signalosome Assembly by DEP Domain Swapping of Dishevelled. Mol Cell 64: 92–104

Hageman JHJ, Keijer J, Dalsgaard TK, Zeper L, Nieuwenhuizen AG (2019) Free fatty acid release from vegetable and bovine milk fat-based infant formulas and human milk during two-phase in vitro digestion. Food & Function 10

Haywood EE, Ho M, Wilson BA (2018) Modular domain swapping among the bacterial cytotoxic necrotizing factor (CNF) family for efficient cargo delivery into mammalian cells. J Biol Chem 293: 3860–3870

Jeffcoat R,., Pollard MR (1977) Studies on the inhibition of the desaturases by cyclopropenoid fatty acids. Lipids 12: 480–485

Kaye Y, Grundman O, Leu S, Zarka A, Zorin B, Didi-Cohen S, Khozin-Goldberg I, Boussiba S (2015) Metabolic engineering toward enhanced LC-PUFA biosynthesis in Nannochloropsis oceanica: Overexpression of endogenous Δ12 desaturase driven by stress-inducible promoter leads to enhanced deposition of polyunsaturated fatty acids in TAG. Algal Research 11: 387–398

Kazuhiro S, Kazuya M, Mikio K, Iwane S, Yasushi T, Koichiro K, Yoshitomo T, Koji M, Masumi H, Naomi K (2004) Functional expression of a Delta12 fatty acid desaturase gene from spinach in transgenic pigs. Proceedings of the National Academy of Sciences of the United States of America 101: 6361–6366

Kikukawa H, Sakuradani E, Kishino S, Park SB, Ando A, Shima J, Ochiai M, Shimizu S, Ogawa J (2013) Characterization of a trifunctional fatty acid desaturase from oleaginous filamentous fungus Mortierella alpina 1S-4 using a yeast expression system. Journal of Bioscience & Bioengineering 116: 672–676

Lamers D, Visscher B, Weusthuis RA, Francke C, Wijffels RH, Lokman C (2019) Overexpression of delta-12 desaturase in the yeast Schwanniomyces occidentalis enhances the production of linoleic acid. Bioresour Technol 289: 121672

Lee JM, Lee H, Kang SB, Park WJ (2016) Fatty Acid Desaturases, Polyunsaturated Fatty Acid Regulation, and Biotechnological Advances. Nutrients 8: 23

Lee K-R, Lee, Yongjik, Kim, Eun-Ha,Lee, Seul-Bee,Roh, Kyung Hee, Kim, Jong-Bum, Kang, Han-Chul, Kim, Hyun Uk (2016) Functional identification of oleate 12-desaturase and ω-3 fatty acid desaturase genes from *Perilla frutescens* var. *frutescens*. Plant cell reports 35: 2523–2537

Liu L, Byeon IJ, Bahar I, Gronenborn AM (2012) Domain swapping proceeds via complete unfolding: a 19F- and 1H-NMR study of the Cyanovirin-N protein. J Am Chem Soc 134: 4229–4235

Luo X, Shi H, Wu R, Wu J, Pi Y, Zheng Y, Yue X (2019) Delta12 fatty acid desaturase gene from Geotrichum candidum in cheese: molecular cloning and functional characterization. FEBS Open Bio 9: 18–25

Marcin G, Sears AE, Kiser PD, Krzysztof P (2015) LRAT-specific domain facilitates vitamin A metabolism by domain swapping in HRASLS3. Nature Chemical Biology 11: 26–32

Meesapyodsuk D, Qiu X (2014) Structure determinants for the substrate specificity of acyl-CoA Delta9 desaturases from a marine copepod. ACS Chem Biol 9: 922–934

Park CK, Joshi HK, Agrawal A, Ghare MI, Little EJ, Dunten PW, Bitinaite J, Horton NC (2010) Domain swapping in allosteric modulation of DNA specificity. PLoS Biol 8: e1000554

Pedrono F, Boulier-Monthean N, Boissel F, Ossemond J, Lohezic-Le Devehat F (2018) The Hypotriglyceridemic Effect of Sciadonic Acid is Mediated by the Inhibition of Delta9-Desaturase Expression and Activity. Mol Nutr Food Res 62

Qin B, Matsuda Y, Mori T, Okada M, Quan Z, Mitsuhashi T, Wakimoto T, Abe I (2016) An Unusual Chimeric Diterpene Synthase from Emericella variecolor and Its Functional Conversion into a Sesterterpene Synthase by Domain Swapping. Angew Chem Int Ed Engl 55: 1658–1661

Ralston JC, Badoud F, Cattrysse B, McNicholas PD, Mutch DM (2014) Inhibition of stearoyl-CoA desaturase-1 in differentiating 3T3-L1 preadipocytes upregulates elongase 6 and downregulates genes affecting triacylglycerol synthesis. Int J Obes (Lond) 38: 1449–1456

Rodríguez-Rodríguez MF, Salas, Joaquín J,Venegas-Calerón, Mónica,Garcés, Rafael,Martínez-Force, Enrique (2016) Molecular cloning and characterization of the genes encoding a microsomal oleate Δ 12 desaturase (CsFAD2) and linoleate Δ 15 desaturase (CsFAD3) from *Camelina sativa*. Industrial Crops and Products 89: 405–415

Sakamoto T, Sakuradani E, Okuda T, Kikukawa H, Ando A, Kishino S, Izumi Y, Bamba T, Shima J, Ogawa J (2017) Metabolic engineering of oleaginous fungus Mortierella alpina for high production of oleic and linoleic acids. Bioresour Technol 245: 1610–1615

ShanlinYu, Pan L, Yang Q, Ping M (2008) Comparison of the Δ12 fatty acid desaturase gene between high-oleic and normal-oleic peanut genotypes. Journal of Genetics & Genomics 35: 679–685

Shi H, Chen H, Gu Z, Song Y, Zhang H, Chen W, Chen YQ (2015) Molecular mechanism of substrate specificity for delta 6 desaturase from *Mortierella alpina* and *Micromonas pusilla*. J Lipid Res 56: 2309–2321

Shi H, Chen H, Gu Z, Zhang H, Chen W, Chen YQ (2016) Application of a delta-6 desaturase with α-linolenic acid preference on eicosapentaenoic acid production in *Mortierella alpina*. Microbial Cell Factories 15: 117

Shi H, Wu R, Zheng Y, Yue X (2018a) Molecular mechanisms underlying catalytic activity of delta 6 desaturase from *Glossomastix chrysoplasta* and *Thalassiosira pseudonana*. J Lipid Res 59: 79–88

Shi H, Xue L, Wu R, Yue X (2018b) Production of eicosapentaenoic acid by application of a delta-6 desaturase with the highest ALA catalytic activity in algae. Microbial cell factories 17: 7

Sun R, Gao L, Yu X, Zheng Y, Li D, Wang X (2016) Identification of a Δ12 fatty acid desaturase from oil palm (Elaeis guineensis Jacq.) involved in the biosynthesis of linoleic acid by heterologous expression in Saccharomyces cerevisiae. Gene 591: 21–26

Sun Z, El Omari K, Sun X, Ilca SL, Kotecha A, Stuart DI, Poranen MM, Huiskonen JT (2017) Double-stranded RNA virus outer shell assembly by bona fide domain-swapping. Nat Commun 8: 14814

Vendome J, Posy S, Jin X, Bahna F, Ahlsen G, Shapiro L, Honig B (2011) Molecular design principles underlying beta-strand swapping in the adhesive dimerization of cadherins. Nat Struct Mol Biol 18: 693–700

Wang H, Klein, Michael G., Zou, Hua, Lane, Weston, Snell, Gyorgy, Levin, Irena, Li, Ke, Sang, Bi-Ching (2015) Crystal structure of human stearoyl-coenzyme A desaturase in complex with substrate. Nature Structural & Molecular Biology 22: 581–585

Wang Y, Zhang S, Pötter M, Sun W, Li L, Yang X, Xiang J, Zhao ZK (2016) Overexpression of Δ12-Fatty Acid Desaturase in the Oleaginous Yeast Rhodosporidium toruloides for Production of Linoleic Acid-Rich Lipids. Applied Biochemistry & Biotechnology 180: 1497–1507

Watanabe K, Oura T, Sakai H, Kajiwara S (2004) Yeast Delta 12 fatty acid desaturase: gene cloning, expression, and function. Journal of the Agricultural Chemical Society of Japan 68: 721–727

Yan Z, Zhuo L, Jiang M, Xia W, Yangmin G, Zhang Y, Huang F (2013) Clone and identification of bifunctional Δ12/Δ15 fatty acid desaturase LKFAD15 from Lipomyces kononenkoae. Food Science & Biotechnology 22: 573–576

Zhang, Chen, Yong Q, Chen, Wei, Chen, Haiqin, Li, Min (2013) Genetic engineering of Yarrowia lipolytica for enhanced production of;trans-10, cis-12 conjugated linoleic acid. Microbial cell factories 12: 70

Zhang Y, Kastman EK, Guasto JS, Wolfe BE (2018) Fungal networks shape dynamics of bacterial dispersal and community assembly in cheese rind microbiomes. Nat Commun 9: 336

Zhang Y, Luan X, Zhang H, Garre V, Song Y, Ratledge C (2017) Improved γ-linolenic acid production in Mucor circinelloides by homologous overexpressing of delta-12 and delta-6 desaturases. Microbial cell factories 16: 113

